# Electrical Stimulation Directs Articular Chondrocyte and Chondrosarcoma Migration in a 3D Collagen Matrix

**DOI:** 10.1101/2025.03.18.643813

**Authors:** J Bush, X Palmer, M Stacey

## Abstract

Electrical signals are fundamental regulators of cell migration and growing numbers of studies have demonstrated electrically guided cancer cell migration. Chondrosarcomas, cartilage forming tumors, are highly metastatic and resistant to chemo and radiation therapies. To measure cellular migration in a three-dimensional (3D) environment a device was 3D-printed to house a collagen gel with embedded cells while enabling direct current electric field (DC-EF) application. Articular chondrocytes and chondrosarcoma cells were exposed to a 1 V/cm electric field for a duration of 12 hours while tracking their migration behavior. We observed that both cell types migrated towards the anode while chondrosarcoma cells showed a stronger directional response. We observed an EF-induced shift from diffusive migration trajectories towards ballistic migration behavior in articular chondrocytes and directed ‘wobble’ migration in chondrosarcoma. In articular chondrocytes we observed significant increases in the length of protrusions directed towards the anode (p < 0.05), as opposed to cathode directed protrusions after EF-exposure. Notably chondrosarcoma cells exhibited tiny protrusions of less than a few microns in length which sporadically extruded and retracted. Chondrosarcoma cells were loaded with FluoVolt to track real-time changes in membrane potential. Cells exposed to a 1V/cm electric field for 30 seconds showed a dynamic cell membrane hyperpolarization and repolarization during EF-exposure with a maximum hyperpolarization approximated to be on the order of −5 mV. To our knowledge, these are the first descriptions of the effects of electrical fields on directional cell migration in a 3D environment.

## INTRODUCTION

Cell migration is a complex phenomenon that can be broken down into key components including cell polarization,^18^ pseudopod formation,^21, 22^ substratum adhesion, cytoskeletal dynamics,^4^ and asymmetric cell contraction. Chondrocytes, the cartilage forming cells, provide an interesting model in which to investigate cell migration. Long considered non-migratory, encased in collagen extra cellular matrix, they contrast widely to the cartilage forming tumors, chondrosarcomas, that are highly metastatic, and are refractory to both chemo-and-radiation therapies. Interestingly, chondrocytes have a relatively depolarized membrane potential (V_m_) like those of undifferentiated, more migratory cell types, including cancer cells. In this study we hypothesized that collagen-embedded human primary chondrocytes and chondrosarcoma cells would migrate directionally in response to an electric field (EF) and that dynamic changes in V_m_ will occur. Reports have shown directional chondrocyte migration when stimulated by electric fields,^7^ ultrasound,^16^ chemotactic mediators,^23^ and enzymatically induced extracellular matrix (ECM) damage.^29^ Chondrocytes’ response to an electric field is well demonstrated and clinical studies have shown superior meniscus healing following their applications.^31^ An accelerated repair of non-articular hyaline cartilage was observed in rats, with a significant increase in cell numbers in the EF treated group.^10, 32^ Furthermore, applied electrical fields impacting membrane potential, may enhance motility and galvanotaxis by affecting intra/extracellular microdomains of ion concentrations. Chondrosarcoma is a metastatic tumor characterized by the formation of a cartilage-like ECM. Among the various steps in the metastatic cascade, it is well-established that cell migration is tightly regulated by the movement of ions.^28^ V_m_ is regarded as an indirect factor that can affect cell migration and the Potassium Inwardly-Rectifying Channel/Gene, Subfamily J, Member 15 (Kir4.2), expressed from the gene *KCNJ15*, has been described as a ‘sensor’ of EFs in tumor cell lines of epithelial and neuronal origin.^24^ Chondrocytes express a rich complement of ion channels which appear to be crucial in the rapidly changing electro-physio environment chondrocytes experience.^1^ In this study we demonstrate that collagen-embedded human primary chondrocytes and chondrosarcoma cells migrate directionally in response to an EF, and that dynamic changes in V_m_ occur. Understanding electric conditions that affect chondrosarcoma migration may be key in opening new avenues of therapeutic research for this aggressive cancer.

## MATERIALS AND METHODS

### Cell Culture

Articular chondrocytes were purchased from PromoCell (Heidelberg Germany) and the chondrosarcoma line JJ012 was kindly donated by Dr. J Block (Rush University Medical Center, Chicago, IL USA). All cells were cultured in Dulbecco’s Modified Eagle Medium (DMEM), (Lonza, MD USA) with 10% Fetal Bovine Serum (Fisher Scientific PA, USA) and 10% Penicillin/Streptomycin (Fisher Scientific PA, USA).

### Device Fabrication

A device was designed with a cell culture volume of 40 mm^3^ for a collagen gel to be embedded during EF application. The device skeleton was 3D printed in a commonly used thermoplastic polymer, Acrylonitrile Butadiene Styrene (ABS). A 10×10mm cover glass was sealed to the upper portion of the chamber with SI-3140 (Dow Corning), a standard glass slide was sealed to the bottom of the 3D-printed chamber (Figure 1). The silicone sealed device was allowed to cure overnight and stored for up to 2 weeks with a 30-minute sterilization under UV light prior to use.

**Figure 1.**
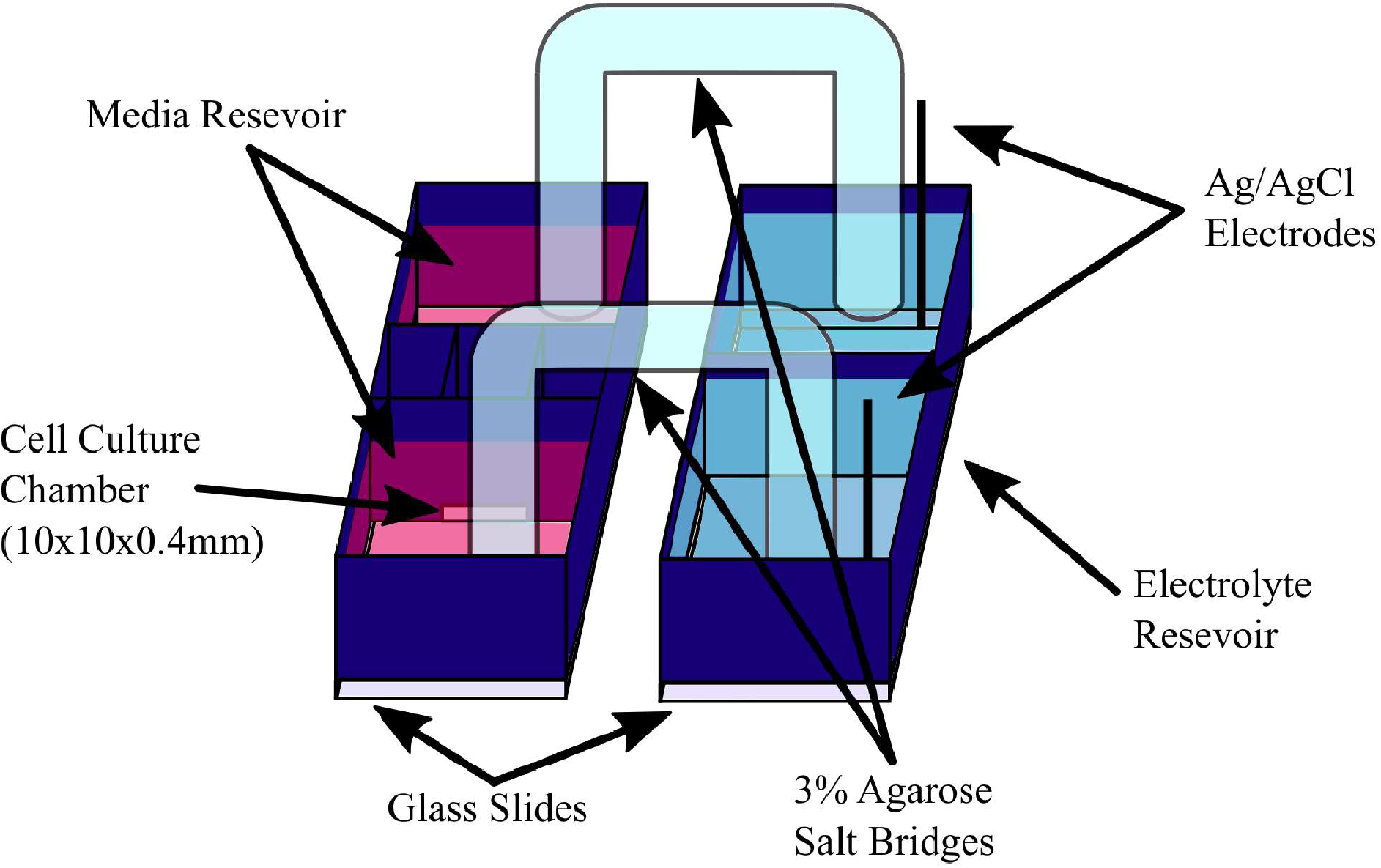
Schematic of the 3D-printed electrotaxis chamber and setup. The electrotaxis chamber contains a 40mm^3^ channel for 3D cell culture and two reservoirs for media. The other device contains two reservoirs filled with Steinberg’s solution and connected to the power supply via Ag/AgCl electrodes measuring 15 cm in length with an inner diameter of 7 mm. The chambers are linked by two glass agar salt bridges. This setup was placed on an ITO heater and housed in a CO_2_ regulated chamber for experiments.

### Collagen Gel and Cell Plating

A 1.5 mg/mL collagen gel mixture was prepared on ice, using Rat Tail Collagen I (Dow Corning). The gel was polymerized at 4 ^o^C for 1 hour to increase the fiber thickness and pore size^11^ permitting adequate cell migration for tracking. Cells were prepared just before the end of this hour. Cells were added to the gel mixture at 90 cells/μL and 40 μL of cell-laden gel solution was plated into the device chamber. The plated gel mixture with embedded cells was polymerized at 37 ^o^C for 30 minutes. Culture media described earlier was then added to the wells on each side of the chamber (Figure 1). Cells were acclimatized in the device for at least 12 hours before experiments.

### Electric Field Application

To apply an electric field to the cells in the device chamber, external solutions of Steinberg’s Solution were interfaced with Ag/AgCl electrodes and Agar Salt Bridges (Figure 1). The silver electrode was chlorinated by soaking for 30 minutes in Clorox Bleach and rinsed with deionized water. The Agar salt bridges were made from 15 cm long glass tubing with an inner diameter of 7 mm. A blow torch was used to soften the hollow glass tubes and form U-shaped glass bridges, which were then sterilized by autoclave, and stored until use. During experimental setup, agarose was mixed with Steinberg’s solution, heated, and inserted into the tubes to form 3% agarose salt bridges. During experiments, cells were maintained at 37 ^o^C, via an Indium Tin Oxide (ITO) heater, and incubated with 5% CO_2_. A voltmeter with stainless steel probe tips was used to measure the voltage across the 1 cm long chamber.

### Image Acquisition and Cell Tracking

For cell migration experiments, images were captured at 2-minute intervals over a 12-hour period using an Olympus IX71 microscope with bright field illumination. Images were aligned using ImageJ mention plug-in. The image stack was saved as a single TIFF file for import into CellTracker, a MATLAB based cell tracking software. CellTracker’s Semi-Automatic Tracking function was used which requires the manual identification of each cell in the initial image, then the migration path is tracked automatically by the software. The raw position coordinates were imported into MATLAB for analysis of the angle of displacement relative to the applied EF, and the Mean-Squared-Displacement (MSD).

### Migration Quantification

The directedness of cell migration was quantified as cos(Φ) where Φ represents the angle between the EF field lines and the resulting vector of the cell’s net displacement. To further quantify the extent of directional cell migration, the MSD of each cell was computed and fitted to the Power Law using the ‘fit’ function in MATLAB, with the 95% confidence interval representing the quality of the fit. The exponent β from the power law fit of the MSD, draws a parallel from diffusive particle motion to provide a quantifiable measure of the cells’ migration trajectories, where a value less than 1 indicates sub-diffusive or ‘wobble’ behavior, a value of 1 indicates purely diffusive behavior, and 2 indicates purely ballistic motion.

### Measurement of Membrane Potential

Cells were loaded with FluoVolt (Thermo Fisher), an electric potential sensitive fluorescent dye, was used to track real-time changes in membrane potential. The dye was loaded into the gel mixture during the gel polymerization step, incubated for 30 minutes, then fresh media was added to the wells as described earlier. Cells were imaged at 1 second intervals for 30 seconds before, during after EF application for a total of 90 seconds. A MATLAB script was written to identify the periphery of the membrane of each cell over time and further segment this into 30 regions based on the angle from the cell’s centroid. To account for photo-bleaching and uneven dye loading, the change in fluorescence was normalized and fitted to a double exponential fit for each segmented region and the mean change in fluorescence reported.

### Statistical Analysis

Statistical analysis was performed using Student t-test to determine significance between the populations, for all tests p<0.05 indicated significance.

## RESULTS

### Chondrocytes migrate towards the anode in a 3D collagen gel under an EF

Articular chondrocytes and chondrosarcoma cells were embedded within a collagen gel and exposed to a DC electric field at 1V/cm. Cell migration was tracked over a period of 12 hours for the control and EF-exposed cells. The directionality of the cells’ migration was quantified as cos(Φ), where Φ represent the angle formed between the electric field lines and an individual cell’s displacement vector. A cos(Φ) value of −1 indicates migration towards the anode parallel to the field lines while a value of 1 indicates same, but towards the cathode. The non-EF-exposed cells showed no directional movement with a directionality of 0.061±0.061 and 0.061±0.0040 for articular chondrocytes and chondrosarcoma cells, respectively. Both cell types responded to the electric field with anodal migration. Articular chondrocytes showed a lower degree of directionality in comparison with chondrosarcoma cells as classified by a mean directionality of −0.47±0.038 and −0.73±0.050, respectively (Figure 2). Statistical analysis yielded p<0.0001 relative to the respective controls for both cell types. Interestingly, the articular chondrocytes exhibited a broader distribution of anode directed migration in comparison with chondrosarcoma cells (Figure 2).

**Figure 2.**
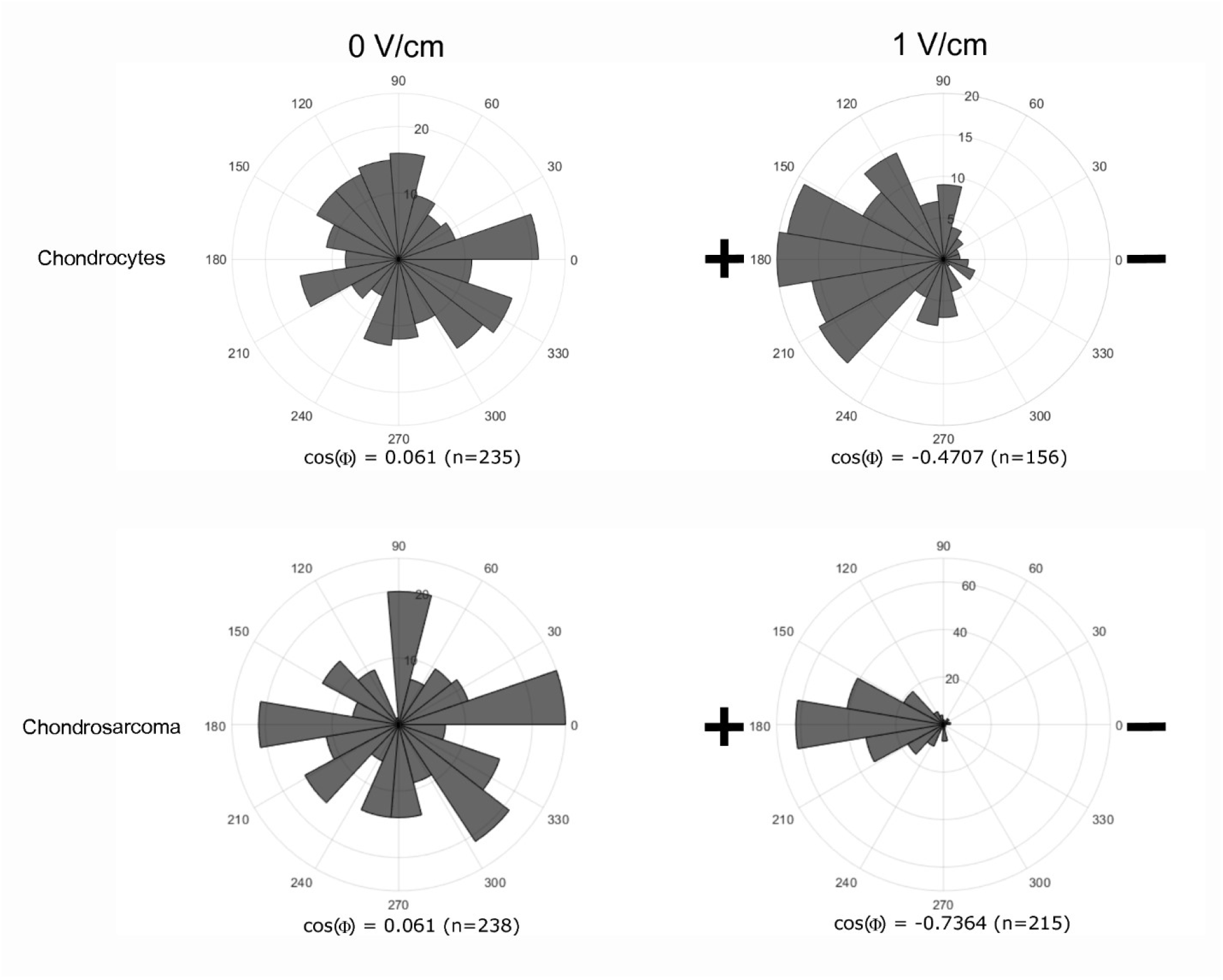
Polar plots showing the cell count and direction of migration for chondrocytes and chondrosarcoma cells when exposed to an EF and under no EF application. The radial axis indicates the number of cells that migrated in a specific direction.

### An external electric field enhances ballistic and persistent cell movement

Both cell types exhibited unique migration patterns within the collagen gel and thus the cos(Φ) metric alone does not necessarily provide an accurate metric with regards to the degree of the cells’ response to the applied field. Specifically, articular chondrocytes migrated with greater persistence and less sporadic protrusion activity, which permitted a greater net displacement, in comparison with the chondrosarcoma cells which displayed tiny but very active and abundant sporadic protrusions that were randomly oriented which resulted in little net displacement, or ‘wobble’ behavior. Wobble behavior results from the sporadic protrusion behavior in conjunction with the gel entrapped state and thus the cos(Φ) metric can be deceptive when quantifying the directedness of migration. For example, a cell may migrate a few microns around its original location during the course of the experiment and have a final position in either direction along the field lines which may yield a cos(Φ) value of ±1, giving the illusion of a stronger directional response when compared to cells migrating with a diffusive trajectory. Thus, we opted to study the cell migration behavior by an additional metric, fitting of the cells’ MSD to the power law of form, ατ^β^. Where α is a scaling factor defined by the net cell displacement, τ is the observed time-step, and β is the power scaling factor, which depicts the dynamics of migration. Commonly used to study anomalous diffusion, an exponent (β) closer to 0, 1, or 2 represents dominant sub-diffusive, diffusive, or ballistic trajectories, respectively. An increase in β under EF exposure indicates an increase in directional migration. Figure 3 shows the average MSD vs τ for both cell types with and without EF application. The function, ατ^β^ was fitted to this data to determine the relevant exponent β, with the 95% confidence interval as the measure of sample deviation. When no electric field is applied, and exponent β equal to 1.09 (95% CI) and 0.605 (95% CI) is obtained for articular chondrocytes and chondrosarcoma cells, respectively (Figure 3). This represents the natural diffusive behavior of unstimulated chondrocytes and confirms the wobble behavior observed visually for chondrosarcoma cells. When a field of 1V/cm is applied an increase in the exponent β was observed from 1.09 (95% CI) to 1.27 (95% CI) for articular chondrocytes and 0.605 (95% CI) to 1.34 (95% CI) for chondrosarcoma cells, confirming a signifcant EF-induced directional response for both cell types.

**Figure 3.**
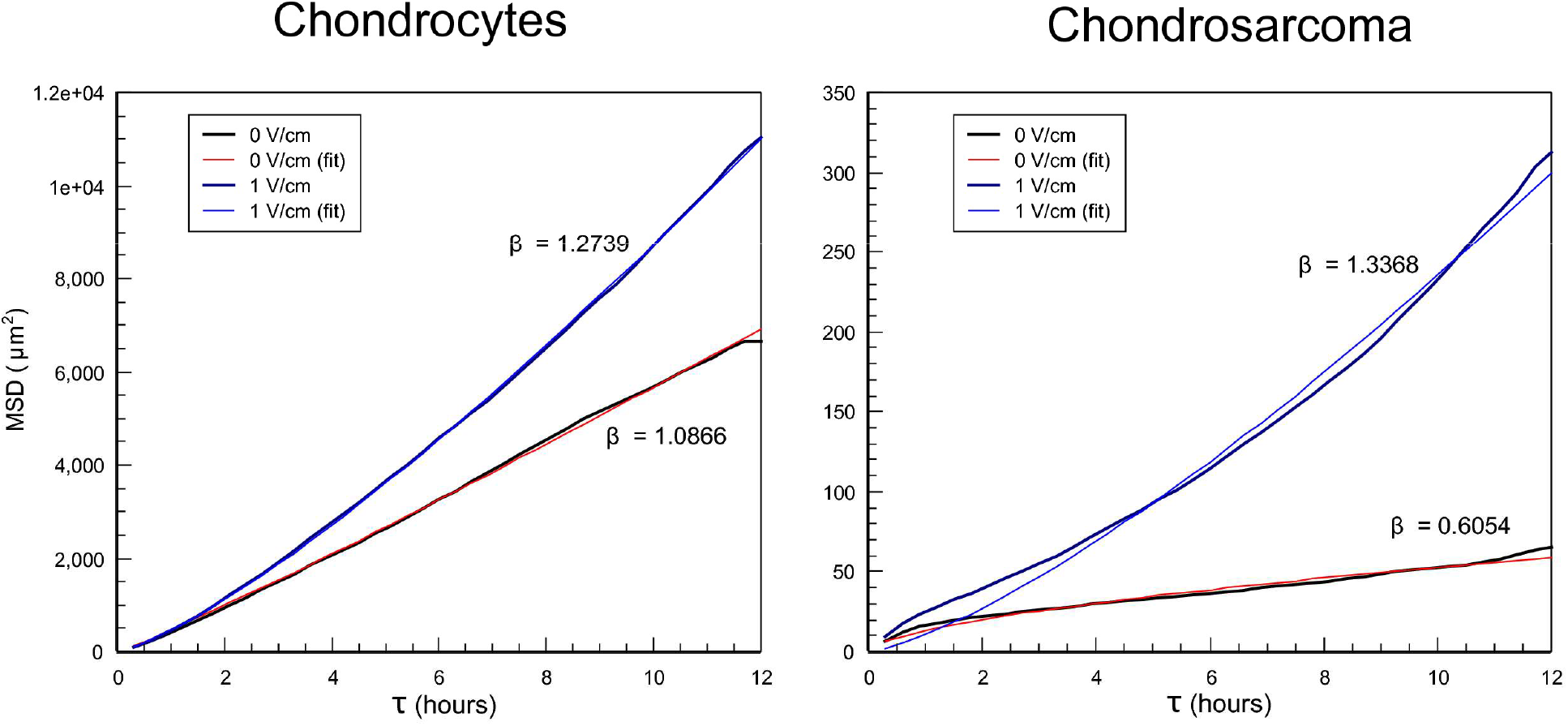
Plots of the average and fitted MSD vs. τ for control and EF-exposed chondrocyte and chondrosarcoma cells. The fitted exponent, β, of each curve fit is labeled next to the respective plot.

### An external electric field enhances the length of anodally directed protrusions

EFs are known to induce asymmetric calcium flux which can play a critical role in the formation of protrusions in migrating cells. Thus, we examined the length of protrusions as measured from the furthest extension to the base at the cell periphery with respect to which the orientation was determined (Figure 4) in unexposed articular chondrocytes and those exposed to 1V/cm for 12 hours. Cells with visible protrusions were identified and cells were selected at random for each condition. In unexposed cells the protrusions showed no specific orientation after 12 hours in culture (p > 0.6) and no significant difference in length (p > 0.6). In chondrocytes exposed to 1V/cm for 12 hours there was no significant difference in the orientation of protrusions (p > 0.4). There was also no significant increase in length of anode or cathode directed protrusions after EF-exposure. However, there was a significant difference in the asymmetric length of protrusions directed towards the anode relative to those directed towards the cathode after EF-exposure (p < 0.05) (Figure 4). Additionally, there was no significant difference in net protrusion length of exposed versus unexposed cells (p > 0.3) or in orientation (p > 0.9). We also examined chondrosarcoma protrusions, which showed many hundreds of small protrusions in all directions, under all conditions, and proved to be unreliable to measure accurately. This observation may be accounted for by the membrane composition of malignant cells. It is known to differ from normal cells where cells with high metastatic potential have lower cholesterol/phospholipid ratios and increased membrane fluidity modulating the structural features of the membrane (viscosity, curvature, electrostatic charge of the bilayer, and migratory potential).

**Figure 4.**
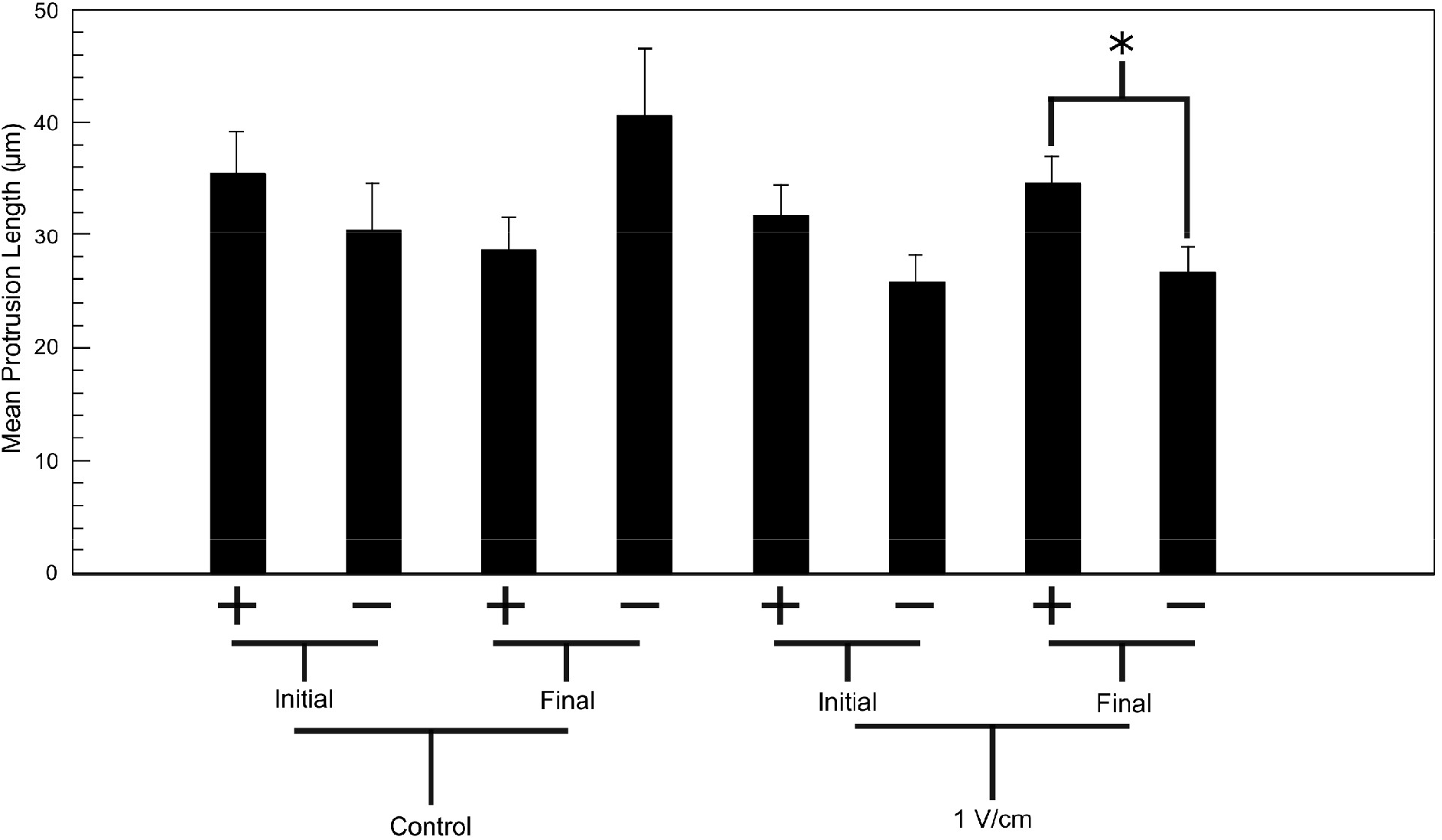
The mean initial and final anode/cathode directed protrusion lengths of articular chondrocytes for control and EF-exposed cells. The +/-indicates anode and cathode directed protrusions respectively. A significant difference (p<0.05) was found only between anode/cathode directed protrusions after EF exposure.

### Dynamic membrane potential response is induced by a 1V/cm electric field

Chondrosarcoma cells were exposed to a 1V/cm electric field for 30 seconds and the fluctuations in membrane potential were reported by the potential-sensitive fluorescent dye, Fluovolt (Thermo Fisher). The membrane potential was examined around the periphery of the cell membrane where dye fluctuations were most detectable (Figure 5). The change in fluorescence of the EF exposed and control cells were compared at individual time points via paired student’s t-test to determine when there was a significant change in membrane potential, before, during and after EF-exposure. Cells that were not exposed to an EF show a stable membrane potential (Fig. 5b), while EF-exposed cells show a transient hyperpolarization, indicated by the decrease in fluorescence, and recovery during EF exposure (Fig. 5c). The EF exposed cells were analyzed in comparison to the control condition at each time point. The time points in where there was a significant difference, (p<0.05) are marked by an asterisk (Fig. 5c). To induce complete depolarization of the cells and confirm a positive response of the dye, cells were loaded with valinomycin, and an isotonic KCl solution was added at t = 30s, (Fig. 5a). Depolarization yielded an average change in fluorescence of 0.20 ± 0.09. The resting membrane potential (RMP) of the chondrosarcoma cell line OUMS-27 is ~ −20 mV,^14^ similar to primary chondrocytes (Lewis et al. 2011). We were not able to identify the RMP of JJ012. Given the previously recorded membrane potential for chondrosarcomas of −20 mV and a relative change in fluorescence (ΔF/F_0_) of 0.20 when cells were depolarized with Valinomycin/KCl, we can approximate the sensitivity of the dye to ~0.01((ΔF/F_0_) / mV) within our setup. Given the maximum change in fluorescence during EF exposure to be −0.047, the magnitude of maximum EF-induced cell hyperpolarization may be approximated as −4.7 mV.

**Figure 5.**
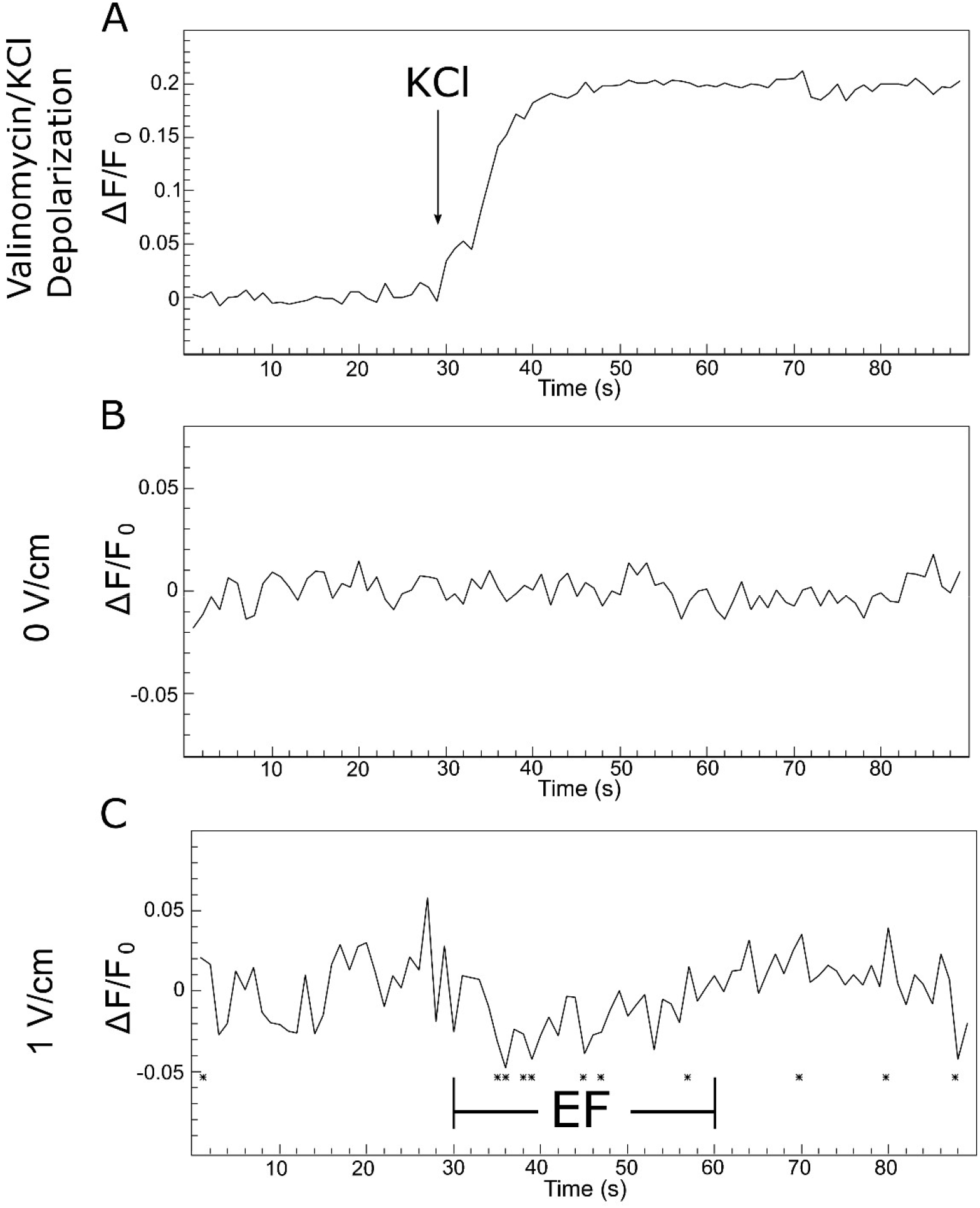
Normalized change in fluorescence vs. time for Fluovolt experiments in which (A) cells were completely depolarized by valinomycin and the addition of KCl (n=10). (B) Change in fluorescence without EF exposure (n=10) and (C) when exposed to an EF for 30 seconds (n=7). Asterisks indicate time points of significant difference (p<0.05) between control and EF-exposed cells.

## DISCUSSION

In this investigation we identified four key features of electrotactic chondrocyte migration in a 3D collagen environment under an EF. Our 3D collagen model is a step towards better resembling the physical environment experienced by these cells compared to 2D glass substrates. It is not however a complete mimic as the gels do not contain the highly negative chondroitin and keratin sulphate proteins that produce the charged environment that gives cartilage many of its properties, for example the high turgidity enabling this tissue to withstand high compressive forces. The electrical sensitivity of chondrocytes questions the description of chondrocytes being essentially passive, contained within a lacuna sensing biomechanical changes within the tissue. Biologically this classic view is incomplete as biological systems are most often integrative. The rich system of cartilage includes heterogeneous mechanical properties, piezoelectric potentials, and proton conductivity due to the presence of structured water on collagen fibers.^13^ It is vital to develop a more holistic knowledge base regarding both the nature of cartilage and how chondrocytes interface with this complex environment.

Most cell types respond to EFs by re-orienting their internal polarity to guide migration, growth, or division. Migrating cells, including epithelial cells, fibroblasts, or neutrophils, respond to EFs by migrating towards the cathode. In contrast, breast cancer cells and some endothelial cells migrate in the opposite direction, toward the anode.^5^ In other cell types, the direction of EF guided cell migration was shown to reverse direction in 2D vs 3D.^15^ To effectively migrate in a 3D matrix, glioma cells are required to degrade the ECM via matrix metalloproteinases and migrate through tortuous confinements with constant osmotically regulated cell deformation and strong retraction forces, but neither of these functions are required in 2D migration.^15^

We show that both articular chondrocytes and a chondrosarcoma cell line (JJ012) migrate towards the anode in a 3D environment. This is contradictory to reports of chondrocytes on 2D glass substrates where migration towards the cathode was observed and notably alignment perpendicular to the field lines was also observed.^7^ It is important to consider that there are many other experimental differences including the origin of the cells, 2D/3D cell morphology, mechanical properties of the substrate, ECM-cell adhesion ligands, and the matrix conductivity. Our results support the significance that environmental factors play in EF-induced chondrocyte polarization and migration responses, thus encouraging progress towards a naturalistic model for the integrative study of electrotactic cues.

EFs signal through the plasma membrane, via ion channels/transporters, to locally activate or recruit components of the cytoskeleton and the polarity machinery. EF sensing has been linked to a variety of mechanisms, including asymmetrical calcium flux, epidermal growth factor receptor activation, and asymmetric transmembrane receptor distribution. Exactly how EFs function to steer polarity remains a challenge to pinpoint at a modeling and molecular level.^5, 6^ Cell migration polarity is integral to a variety of physiological processes including organ development, tissue morphogenesis, wound healing, and immune responses. A greater understanding of the effects environmental cues have on motility can inform the design of biotechnologies such as movement-directing scaffolds.^20^ Typically, migration through a matrix involves the generation of cell protrusions, i.e. extensions of the plasma membrane beyond the cell body, which are driven by actin polymerization and osmotic pressure. Our cell migration data showed that in response to an EF, cell migration became more directed with a shift from diffusive to ballistic trajectory as indicated by the relative angle of displacement (Figure 2) and the MSD (Figure 3). Diffusive migration in 3D is most probable due to the complex interactions between the cell and its surrounding heterogeneous ECM, thus diminishing the effects of external migratory stimuli, such as an EF,^17^ and lowering the potential for true ballistic migration, as commonly seen in 2D. Measuring the number, orientation, and length of protrusions we identified a significant asymmetry in the length of protrusions after an EF was applied, which showed longer protrusion in the direction of the anode, relative to those directed towards the cathode.

Sarcomas have recently been shown to express significant levels of ion channels.^2^ Given the high expression of ion channels, it was of interest to determine whether an electric field could stimulate the cells to dynamically alter their membrane potential. Particularly since calcium flux has been identified as a primary mechanism in the electrotactic migration response of multiple cell lines.^3^ Interestingly the JJ012 chondrosarcoma cell line exhibited a stronger directional response, as quantified by cos(Φ), than the articular chondrocyte cell line. The cohesive role ion channels play in sensing EFs remains to be developed towards a greater understanding. To assess the collective response of ion channels in chondrosarcoma cells exposed to a 1V/cm EF, membrane potential was monitored using the potential sensitive dye, Fluovolt, before, during, and after EF exposure.

A large number of voltage gated ion channels have been reported in chondrocytes by gene array analysis.^1^ Due to the ability of these channels to differentially transfer ions across the cell membrane in response to an electric field, we chose to investigate dynamic EF-induced changes in plasma membrane potential. We observed a significant whole cell hyperpolarization for ~3 seconds after initiating EF exposure which then persisted for ~10-20 seconds whilst being dynamically corrected by the cell during EF exposure and finally settling to baseline after EF exposure (Fig. 5c). The hyperpolarization timescale is consistent with that of weaker calcium spikes induced by calcium injection of neighboring chondrocytes.^9^ EF-induced calcium influx can stimulate intracellular calcium release resulting in a calcium wave and substantial increase in intracellular calcium. However, positively regulated intracellular calcium release would result in depolarization of the cells. It was previously shown that chondrocytes exposed to stronger EFs of 2, 4, 6 V/cm in 2D conditions migrate towards the cathode. Neomycin, an inhibitor of Inositol Triphosphate (IP3) mediated intracellular Ca^2+^ release,^27^ inhibited the cathodal migration response and even caused some cells to migrate anodally, while EFs < 2V/cm elicited no significant directional response in 2D conditions.^7^ Stronger EFs may induce a greater calcium influx thus overcoming the threshold for positively regulated intracellular calcium release. Under an EF of lower magnitude, localized calcium influx may play the dominant role in directing migration. Concentrated microdomains of calcium influx can develop near calcium activated potassium channels as Large-Conductance, Voltage, and Calcium-Activated Potassium Channels (BK) and Small-Conductance Calcium-Activated Potassium Channels (SK2) channels have been shown to colocalize with voltage gated calcium channels.^19, 30^ Calcium-activated potassium channels such as KCa3.1 have been shown to play a significant role in cell migration, though the complete mechanism remains unclear.^8, 12^

## CONCLUSION

Our investigation showed that both articular chondrocytes and chondrosarcoma cells respond to a physiologically relevant electric field of 1 V/cm. Both chondrocytes and chondrosarcoma cells respond with anodally directed migration and chondrocytes showed enhanced protrusion length towards the anode. To begin probing the ion channel related mechanisms for EF sensing in collagen embedded chondro-related cells, we studied the response of chondrosarcoma membrane potential to electric fields and found a transient hyperpolarization and stabilization during EF exposure, thus confirming a significant response of ion channels. These results provide a macroscopic view of EF modulated ion flux to help direct future work which may focus on specific ion conduction cascades and their role in the electrotactic migration response of chondrocytes and chondrosarcoma cells. As well, since chondrocytes regularly exist in clusters within the cartilage, it will be crucial to further explore their intercellular response to electric fields, particularly their role in regenerative processes.

## ACKNOWLEDGEMENT

This work was partially funded by a grant from the Breeden Adams Foundation.

## REFERENCES

1. Asmar, A., R. Barrett-Jolley, A. Werner, R. Kelly, Jr. and M. Stacey (2016). “Membrane channel gene expression in human costal and articular chondrocytes.” Organogenesis 12(2): 94–107.

2. Aung, T., C. Asam and S. Haerteis (2019). “Ion channels in sarcoma: pathophysiology and treatment options.” Pflugers Arch 471(9): 1163–1171.

3. Babona-Pilipos, R., N. Liu, A. Pritchard-Oh, A. Mok, D. Badawi, M. R. Popovic and C. M. Morshead (2018). “Calcium influx differentially regulates migration velocity and directedness in response to electric field application.” Exp Cell Res 368(2): 202–214.

4. Blain, E. J. (2009). “Involvement of the cytoskeletal elements in articular cartilage homeostasis and pathology.” Int J Exp Pathol 90(1): 1–15.

5. Bonazzi, D. and N. Minc (2014). “Dissecting the Molecular Mechanisms of Electrotactic Effects.” Adv Wound Care (New Rochelle) 3(2): 139–148.

6. Campetelli, A., D. Bonazzi and N. Minc (2012). “Electrochemical regulation of cell polarity and the cytoskeleton.” Cytoskeleton (Hoboken) 69(9): 601–612.

7. Chao, P. H., R. Roy, R. L. Mauck, W. Liu, W. B. Valhmu and C. T. Hung (2000). “Chondrocyte translocation response to direct current electric fields.” J Biomech Eng 122(3): 261–267.

8. Cruse, G., S. M. Duffy, C. E. Brightling and P. Bradding (2006). “Functional KCa3.1 K+ channels are required for human lung mast cell migration.” Thorax 61(10): 880–885.

9. D’Andrea, P. and F. Vittur (1996). “Ca2+ oscillations and intercellular Ca2+ waves in ATP-stimulated articular chondrocytes.” J Bone Miner Res 11(7): 946–954.

10. de Campos Ciccone, C., D. C. Zuzzi, L. M. Neves, J. S. Mendonca, P. P. Joazeiro and M. A. Esquisatto (2013). “Effects of microcurrent stimulation on hyaline cartilage repair in immature male rats (Rattus norvegicus).” BMC Complement Altern Med 13: 17.

11. Doyle, A. D., N. Carvajal, A. Jin, K. Matsumoto and K. M. Yamada (2015). “Local 3D matrix microenvironment regulates cell migration through spatiotemporal dynamics of contractility-dependent adhesions.” Nat Commun 6: 8720.

12. Duffy, S. M., G. Cruse, S. L. Cockerill, C. E. Brightling and P. Bradding (2008). “Engagement of the EP2 prostanoid receptor closes the K+ channel KCa3.1 in human lung mast cells and attenuates their migration.” Eur J Immunol 38(9): 2548–2556.

13. Friesen, D. E., T. J. Craddock, A. P. Kalra and J. A. Tuszynski (2015). “Biological wires, communication systems, and implications for disease.” Biosystems 127: 14–27.

14. Funabashi, K., M. Fujii, H. Yamamura, S. Ohya and Y. Imaizumi (2010). “Contribution of chloride channel conductance to the regulation of resting membrane potential in chondrocytes.” J Pharmacol Sci 113(1): 94–99.

15. Huang, Y. J., G. Hoffmann, B. Wheeler, P. Schiapparelli, A. Quinones-Hinojosa and P. Searson (2016). “Cellular microenvironment modulates the galvanotaxis of brain tumor initiating cells.” Sci Rep 6: 21583.

16. Jang, K. W., L. Ding, D. Seol, T. H. Lim, J. A. Buckwalter and J. A. Martin (2014). “Low-intensity pulsed ultrasound promotes chondrogenic progenitor cell migration via focal adhesion kinase pathway.” Ultrasound Med Biol 40(6): 1177–1186.

17. Jiang, C., C. Cui, L. Li and Y. Shao (2014). “The anomalous diffusion of a tumor invading with different surrounding tissues.” PLoS One 9(10): e109784.

18. Kuss, P., K. Kraft, J. Stumm, D. Ibrahim, P. Vallecillo-Garcia, S. Mundlos and S. Stricker (2014). “Regulation of cell polarity in the cartilage growth plate and perichondrium of metacarpal elements by HOXD13 and WNT5A.” Dev Biol 385(1): 83–93.

19. Lu, L., P. Sirish, Z. Zhang, R. L. Woltz, N. Li, V. Timofeyev, A. A. Knowlton, X. D. Zhang, E. N. Yamoah and N. Chiamvimonvat (2015). “Regulation of gene transcription by voltage-gated L-type calcium channel, Cav1.3.” J Biol Chem 290(8): 4663–4676.

20. Luzhansky, I. D., A. D. Schwartz, J. D. Cohen, J. P. MacMunn, L. E. Barney, L. E. Jansen and S. R. Peyton (2018). “Anomalously diffusing and persistently migrating cells in 2D and 3D culture environments.” APL Bioeng 2(2): 026112.

21. Lyman, J. R., J. D. Chappell, T. I. Morales, S. S. Kelley and G. M. Lee (2012). “Response of Chondrocytes to Local Mechanical Injury in an Ex Vivo Model.” Cartilage 3(1): 58–69.

22. Mayan, M. D., R. Gago-Fuentes, P. Carpintero-Fernandez, P. Fernandez-Puente, P. Filgueira-Fernandez, N. Goyanes, V. Valiunas, P. R. Brink, G. S. Goldberg and F. J. Blanco (2015). “Articular chondrocyte network mediated by gap junctions: role in metabolic cartilage homeostasis.” Ann Rheum Dis 74(1): 275–284.

23. Mishima, Y. and M. Lotz (2008). “Chemotaxis of human articular chondrocytes and mesenchymal stem cells.” J Orthop Res 26(10): 1407–1412.

24. Nakajima, K. I., K. Zhu, Y. H. Sun, B. Hegyi, Q. Zeng, C. J. Murphy, J. V. Small, Y. Chen-Izu, Y. Izumiya, J. M. Penninger and M. Zhao (2015). “KCNJ15/Kir4.2 couples with polyamines to sense weak extracellular electric fields in galvanotaxis.” Nat Commun 6: 8532.

25. Nuccitelli, R. (2003). “A role for endogenous electric fields in wound healing.” Curr Top Dev Biol 58: 1–26.

26. Pietak, A. and M. Levin (2018). “Bioelectrical control of positional information in development and regeneration: A review of conceptual and computational advances.” Prog Biophys Mol Biol 137: 52–68.

27. Prentki, M., J. T. Deeney, F. M. Matschinsky and S. K. Joseph (1986). “Neomycin: a specific drug to study the inositol-phospholipid signalling system?” FEBS Lett 197(1-2): 285–288.

28. Schwab, A., A. Fabian, P. J. Hanley and C. Stock (2012). “Role of ion channels and transporters in cell migration.” Physiol Rev 92(4): 1865–1913.

29. Seol, D., Y. Yu, H. Choe, K. Jang, M. J. Brouillette, H. Zheng, T. H. Lim, J. A. Buckwalter and J. A. Martin (2014). “Effect of short-term enzymatic treatment on cell migration and cartilage regeneration: in vitro organ culture of bovine articular cartilage.” Tissue Eng Part A 20(13-14): 1807–1814.

30. Vierra, N. C., M. Kirmiz, D. van der List, L. F. Santana and J. S. Trimmer (2019). “Kv2.1 mediates spatial and functional coupling of L-type calcium channels and ryanodine receptors in mammalian neurons.” Elife 8.

31. Yuan, X., D. E. Arkonac, P. H. Chao and G. Vunjak-Novakovic (2014). “Electrical stimulation enhances cell migration and integrative repair in the meniscus.” Sci Rep 4: 3674.

32. Zuzzi, D. C., C. Ciccone Cde, L. M. Neves, J. S. Mendonca, P. P. Joazeiro and M. A. Esquisatto (2013). “Evaluation of the effects of electrical stimulation on cartilage repair in adult male rats.” Tissue Cell 45(4): 275–281.

